# Impact of chemotherapy on leukocyte stiffness -A longitudinal RT-DC study in a breast cancer patient

**DOI:** 10.64898/2026.07.07.736962

**Authors:** Martin Kräter, Christoph Herold, Anna Taubenberger, Nicole Töpfner, Marta Urbanska, Maik Herbig, Theresa Link, Martin Bornhäuser, Jochen Guck, Angela Jacobi

## Abstract

**Background:** The physical properties of leukocytes, such as cell size and stiffness, are critical for their circulation in microcapillary networks where rapid shape changes are required to squeeze through vascular constrictions. Alterations in the cells’ physical phenotype can promote venous thromboembolism (VTE), a common cause of death in cancer patients receiving chemotherapy. While biochemical VTE predictors are well studied, physical properties of blood cells receive less attention.

**Methods:** Using real-time deformability cytometry (RT-DC), we monitored for the first time the physical phenotype of leukocytes in a longitudinal study of a breast cancer patient treated with epirubicin/cyclophosphamide (EC) and paclitaxel (Pax).

**Results:** The leukocyte counts extracted from RT-DC were in good agreement with standard clinical leukograms and EC had no immediate effect on leukocyte properties. However, Pax caused a significant softening of granulo/monocytes and a stiffening of lymphocytes immediately after administration. Leukocyte size was constant throughout the therapy, but we observed an overall increase in leukocyte stiffness, which was restored to normal values 45 weeks post treatment.

**Conclusion:** Taken together, our data reveal chemotherapy-induced specific alterations of leukocyte stiffness potentially critical for microcirculation. Thus, RT-DC measurements can add important, yet currently not available information to VTE prediction in cancer patients.

## Background

Chemotherapeutic drugs often inhibit cell growth and division by interfering with the cytoskeleton, for example by arresting mitosis through stabilisation/destabilisation of microtubules. However, in addition to its role in cell division, the cytoskeleton is also a primary determinant of a cell’s physical properties, such as size and stiffness ^1^. While the tumour is the primary target of chemotherapy, less is known about this therapy’s physical effects on leukocytes, which are immediately impacted following venous drug administration.

Leukocyte physiological functions such as immune responses rely on the ability to circulate freely within the blood vessels and transmigrate to target tissues. The efficiency of these processes largely depend on the cytoskeleton and interfering with it can vitally disturb immune function (for review see ^2^) and hamper vascular passage ^3,4^. Previous studies have shown that rapid changes in cell size and stiffness are required for squeezing through vascular constrictions ^5–7^, and alterations in these properties can potentially be used to detect disease conditions by probing blood cells ^8^. Chemotherapeutic treatment was shown to induce stiffening of leukemic cells potentially reducing vascular passage in leukaemia patients ^9^. Subsequently, mechanical changes in blood cells can lead to reduced blood flow (for review see ^10^) potentially initiating venous complications (VCs) such as venous thromboembolism by vascular occlusion. VCs are a leading cause of death in cancer patients receiving outpatient chemotherapy ^11^ and numerous studies seek to identify biochemical markers that can predict patients at different risk levels (for review see ^12^). Although leukocytes are known to contribute for example to thrombosis development after phenotypic transformation (for review see ^13^) and blood counts seem to be an important indicator ^14^, the physical properties of leukocytes have so far received less attention.

Here, we report the first longitudinal study to assess the physical properties of leukocytes of a breast cancer patient undergoing chemotherapy. We employed real-time deformability cytometry (RT-DC), a high-throughput technology that allows label free, image-based morphological and mechanical interrogation of cells at rates of up to 1000 cells per second ^15^. The patient was treated with a standard primary systemic therapy used in breast cancer treatment including administration of epirubicin/cyclophosphamide (EC) followed by paclitaxel (Pax) (**Figure 1A**). EC, an anthracycline-based combination, intercalates into the DNA and interferes with cell replication ^16,17^. Pax promotes microtubule assembly and stabilization, resulting in cell cycle arrest and inhibition of proliferation (for review see ^18^). Anthracyclines have been shown to affect cell circulation *in vitro* ^19^ and Pax likely interferes with the physical properties of leukocytes as it stabilizes a major compartment of the cell’s cytoskeleton (for review see ^18^). One could therefore speculate, that alterations in cell-stiffness may affect leukocyte microcirculation with secondary effects on VC. Motivated by this longitudinal study we envision an RT-DC-based approach for predicting impaired leukocte function and vessel retention using blood cell physical phenotyping.

**Figure 1.**
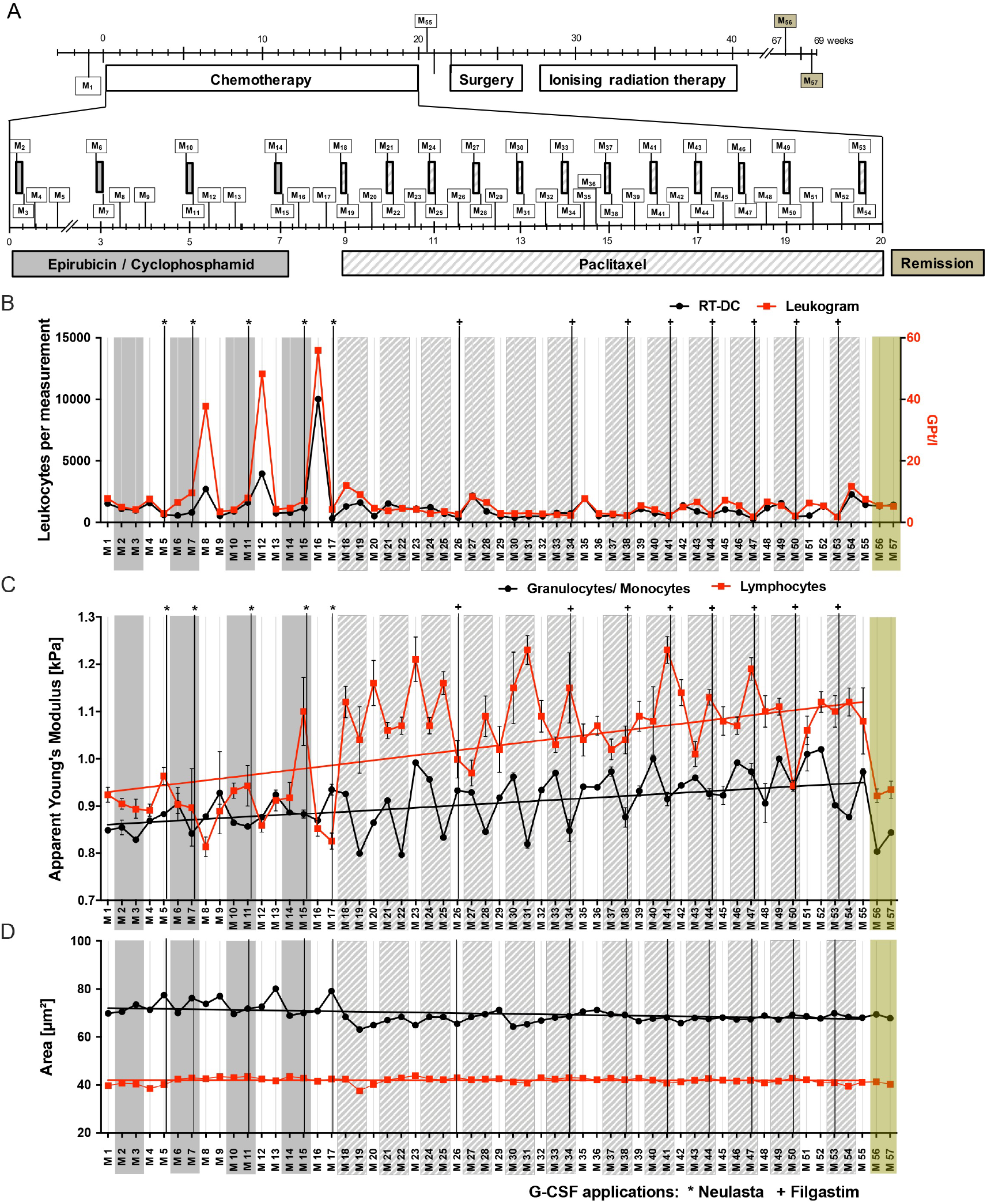
Physical changes of leukocytes during chemotherapy. *(A)* The therapy schedule of a neoadjuvant chemotherapy treated breast cancer patient including four cycles of epirubicin/ cyclophosphamide (E/C) treatment (seven weeks, biweekly treatment) followed by 12 cycles of paclitaxel treatment (Pax) (twelve weeks, weekly treatment). 57 RT-DC blood measurements (M_1-57_) were performed during the course of this study as indicated by rectangles. During the chemotherapy treatment, measurements were done immediately before and after every treatment cycle as well as three days after treatment (white zones between the grey boxes). Additionally, measurements M36 and M52 were carried out one day later because the patient’s therapy schedule had to be postponed by one day for health reasons. *(B)* Leukocyte count over the therapy time course. Administration of G-CSF (*Neulasta or ^+^Filgastrim) is shown by vertical lines. RT-DC-obtained leukocyte count per 15 min measurement is displayed for every measurement as interconnected black dots and compared to a standard clinical leukogram (interconnected red squares). *(C+D)* Apparent Young’s Modulus and cell cross-sectional area of granulo-/ monocytes (interconnected black dots) and lymphocytes (interconnected red squares) obtained from RT-DC measurements throughout therapy. Dots and squares represent the median values and error bars indicate the standard error. The line is only a graphical aid for the time-dependent changes and can be seen as a visual guide. Median values, standard deviations (SDs) and standard error (SEMs) for apparent Young’s modulus (YM), area and deformation for M1-M57 for granulo-/ monocytes can be found in supplement table 1 and for lymphocytes in supplement table 2.

## Materials and Methods

Peripheral blood (PB) was obtained from a 33-year-old female patient after informed consent, in accordance with the Declaration of Helsinki. Ethical approval for experiments with human blood were approved by the ethics committee of the Technische Universität Dresden (EK89032013, EK458102015).

The patient received a neoadjuvant standard therapy with four cycles of epirubicin and cyclophosphamide (EC) followed by twelve cycles of paclitaxel (Pax) over a period of 20 weeks (**Figure 1A**). To counteract low blood cell count, granulocyte colony stimulation factor (G-CSF; Neulasta and Filgastrim) was administered throughout therapy as indicated in **Figure 1A**. Venous blood was drawn with a 20-gauge multifly needle into a sodium citrate tube (S-Monovette® 10 mL 9NC, Sarstedt, Germany) by vacuum aspiration. Measurements were carried out within 2 hours. A conventional whole blood count (XE-5000, Sysmex, Norderstedt, Germany), was performed for every time point (indicated as Leukogram in **Figure 1B**), in parallel with the RT-DC measurement.

Real-time deformability cytometry (RT-DC) was performed as described elsewhere ^8,15,20^. Briefly, 50 µl of anti-coagulated blood was diluted in 950 µl measurement buffer (MB). The MB was based on PBS, supplemented with 0.6% (w/v methyl cellulose, adjusted to a viscosity of 25 ± 0.6 mPa·s at 24°C. The cell suspension was pumped through a microfluidic chip by means of a syringe pump (NemeSyS, Cetoni, Germany) at a flow rate of 0.015 µl/s. A second syringe pump operating at 0.045 µl/s supplied a sheath flow, which focussed the sample flow towards a narrow squared channel (20 µm x 20 µm cross-section) within the microfluidic chip. At the end of the channel, the cell images were captured by a high-speed camera and their contour was determined. From the contour, the cross-sectional area (*A* in µm^2^) and deformation 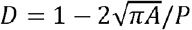, where *P* is the perimeter of the contour) were derived in real-time. *A* and *D* were then used to identify the apparent Young’s modulus of each initially spherical leukocyte, under the assumption of a homogeneous, solid, and linearly elastic object ^21,22^. Every blood sample was measured for 15 minutes resulting in a total leukocyte count per measurement as illustrated in **Figure 1B**.

RT-DC data were analyzed using ShapeOut2 (version: 2.4.15 available at https://github.com/ZELLMECHANIK-DRESDEN/ShapeOut/releases). With a brightfield image for every measured cell, the acquired data allow for multiparametric offline analysis to discriminate between different cell types. Both, lymphocytes and granulo-/ monocytes can be robustly identified and gated by area ratio, cell size and deformation, as described by Toepfner et al. ^8^ and illustrated in **supplement Figure 1A and B**. Statistical analysis was carried out using one-dimensional linear mixed models (LMM) that incorporate fixed effect parameters and random effects, allowing to describe inherent variability between replicated measurements. For computation of significance levels, a likelihood ratio test was employed to compare an LMM, which assumes an effect, with a second LMM, where the effect term is set to zero ^23^.

## Results and Discussion

Over the course of this study, we acquired leukocyte physical features from a total of 57 RT-DC measurements of unaltered whole blood samples (**Figure 1A**). Two measurements were performed before the onset of chemotherapy treatment, followed by 53 measurements during drug administration, and two measurements 45 weeks after the last treatment cycle (remission). The accuracy of RT-DC measurements to detect leukocytes was verified by comparing leukocyte counts of each measurement to corresponding standard leukograms (**Figure 1B**). In accordance with our previous report ^8^, we found that cell counts established with RT-DC correlated with leukogram-based counts but somehow underestimated the standard counts by percent. Upon G-CSF (Neulasta and Filgastim) administration, we observed an increase in leukocyte count both in the leukograms and RT-DC measurements; though it was less pronounced in the RT-DC-derived counts (**Figure 1B**). Interestingly, a trend towards a decrease in leukocyte stiffness after G-CSF administration was observed, most prominent with Neulasta treatment **(Figure 1C)**. In line with this observation, Fay et al. had postulated that a rapid decrease in leukocyte stiffness could be a first step in leukocyte demargination and trafficking, thereby increasing clinical blood counts ^5^.

Throughout the whole therapy, a long-term increase in lymphocyte (∼ 20%) and granulo-/ monocyte (∼ 10%) stiffness was observed **(Figure 1C)**, reflected by the general decrease in deformability measured in RT-DC (**Supplement Figure 1C**). Previously, Lam et al. had reported chemotherapy-induced leukemic cell stiffening correlated with apoptosis induction ^9^. Both effects, stiffening and apoptosis induction, potentially contribute to the observed reduced counts of circulating leukocytes. However, the leukocyte stiffening over the entire therapy period most likely cannot be attributed to apoptosis induction, as with measurements between the therapy cycles we probe already renewed blood cells. This is indicative for an effect of the therapy on the haematopoietic organ, as described for Pax and cyclophosphamide in combination with rhG-CSF administration^24^. Interestingly, leukocyte stiffness had returned to basal levels 45 weeks after the final chemotherapy cycle (**Figure 1C**). The average cell size, quantified as projected cross-sectional area, remained constant throughout the treatment, when compared to the size before chemotherapeutic intervention or during remission period. Only some small fluctuations in cell size were observed during EC administration (**Figure 1D**).

We found a remarkable short-term decrease in granulo-/ monocyte stiffness immediately after Pax treatment. This effect was observed in every Pax cycle (**Figure 2A-C**) and is in line with the finding of Golfier et al., who showed an increased deformation of Pax treated HL-60 cells using RT-DC ^25^. Interestingly, the immediate decrease in stiffness of granulo-/ monocytes was becoming progressively smaller, but still detectable, in sequential treatment cycles (**Figure 2A-C**). This suggests that with progressing chemotherapy treatment the composition and assembly of the cytoskeleton, or its response to the same stimulus was altered. Contrary to the granulo-/monocytes, lymphocytes became stiffer after Pax treatment (**Figure 2 D-F**), which to our knowledge has not been reported yet. Of note, EC treatment showed no short-term effect either on lymphocyte or granulo-/ monocyte properties (**Figure 1C**).

**Figure 2.**
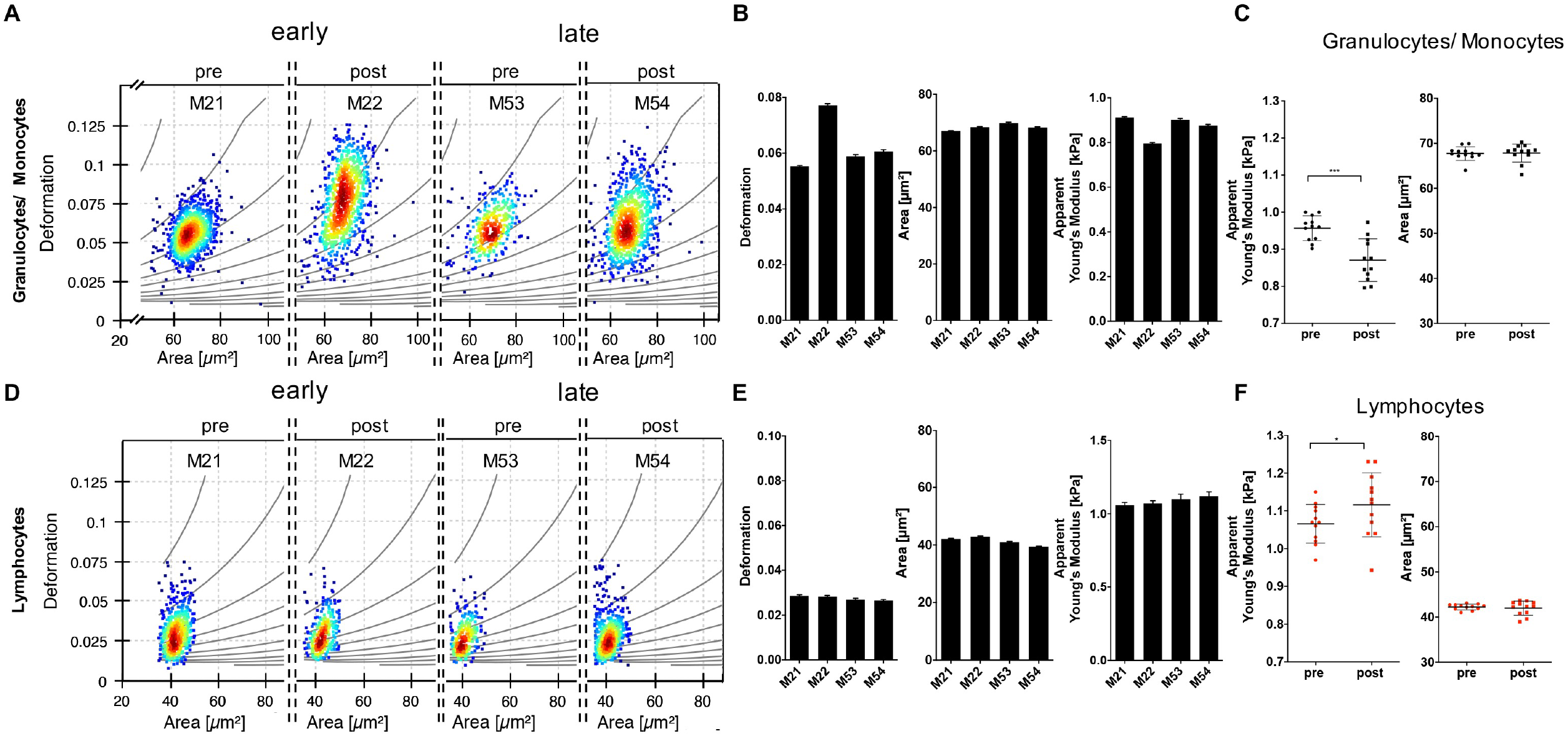
Short-term deformability of leukocytes is increased by paclitaxel treatment. *(A)* Early (M21 & M22) and late (M53 & M54) treatment phase-derived scatter-plots of deformation versus area of granulo-/ monocytes before (M21 & M53) and after (M22 & M54) two representative Pax treatments. Thin, curved lines in the background are isoelasticity lines that indicate grouping according to apparent Young’s modulus (YM). *(B)* Bar-graphs show median values ± SEM of cell deformation, area, and apparent YM of granulo-/ monocytes before (M21 & M53) and after (M22 & M54) two representative Pax treatments. *(C)* Apparent YM in kPa and cross-sectional area values in µm^2^ of granulo-/ monocytes. Diagram represents median values ± SD of 12 Pax cycles pre- and post-treatment. *(D)* Early (M21 & M22) and late (M53 & M54) treatment phase-derived scatter-plots of deformation versus area of lymphocytes before (M21 & M53) and after (M22 & M54) two representative Pax treatments. *(E)* Median values ± SEM of cell deformation, area, and apparent YM of lymphocytes before (M21 & M53) and after (M22 & M54) two representative Pax treatments. *(F)* Median values of apparent YM in kPa and area values in µm^2^ of lymphocytes before and after 12 Pax cycles (pre- and post-treatment). For statistical analysis, an approach using linear mixed models was employed (* *p* = 0.038***; *p* = 0.00031).

## Conclusion

VC-risk is determined at later stages of the disease by various risk factors concerning activation of plasmatic coagulation and endothelial cell activity. However, altered leukocyte mechanical properties may contribute to vascular occlusion and reduced hemodynamic flow (for reviewed see ^26^), finally initiating VCs. Measurements of physical properties of leukocytes during chemotherapy of a large cohort of patients might elicit our understanding of whether VCs are additionally promoted by leukocyte stiffness changes. In addition, changes in leukocyte circulative properties could theoretically affect aspects of innate immunity and jeopardize the defense against infections beyond the consequences of low neutrophil and lymphocyte counts.

In conclusion, we show for the first time in a patient undergoing chemotherapy that EC treatment has no immediate effect on size and stiffness of leukocytes, while Pax treatment transiently softens granulo-/ monocytes and stiffens lymphocytes after administration. We consider this effect direct to the circulating cells. However, EC and Pax administration have an overall long-term stiffening effect on all leukocytes, which potentially indicates an effect of the drugs on the hematopoietic organ or directly on the hematopoietic stem cells. The follow-up measurements 45 weeks after treatment indicate a complete reversion of the long-term stiffening to similar levels observed prior to treatment. Most importantly, we demonstrate that size and stiffness characteristics of blood cells can readily be monitored in patients using RT-DC, potentially providing a potent tool to predict the risk of VTE caused mortality.

## Abbreviations

EC: epirubicin and cyclophosphamide
Pax: paclitaxel
PB: peripheral blood
LMM: linear mixed models
VC: venous complications
VTE: venous thromboembolism
RT-DC: real-time deformability cytometry
G-CSF: granulocyte colony stimulation factor

## Ethics approval and consent to participate

Human subjects: The work involved measurements of human blood samples. All studies complied with the Declaration of Helsinki and involved written informed consent from the participant. Ethics for experiments with human blood were approved by the ethics committee of the Technische Universität Dresden (EK89032013, EK458102015).

## Consent

Written informed consent was obtained from the patient for publication of this publication.

## Availability of data and material

Original raw data are available.

## Acknowledgements

The authors would like to thank the oncological ambulance team for their support as well as Manja Wobus for proofreading the manuscript.

## Authors contributions

A.J. and M.K. designed the project outlined and carried out most experiments, interpreted results and co-wrote the manuscript. A.T., C.H., N.T. and T.L. acquired data, supported study design and interpretation. A.T., M.U., M.H., and D.S. supported data analysis and interpretation. M.B. and J.G. supported interpretation. J.G. and A.J acquired funding. All authors revised and edited the manuscript.

---

**Supplement Figure.**
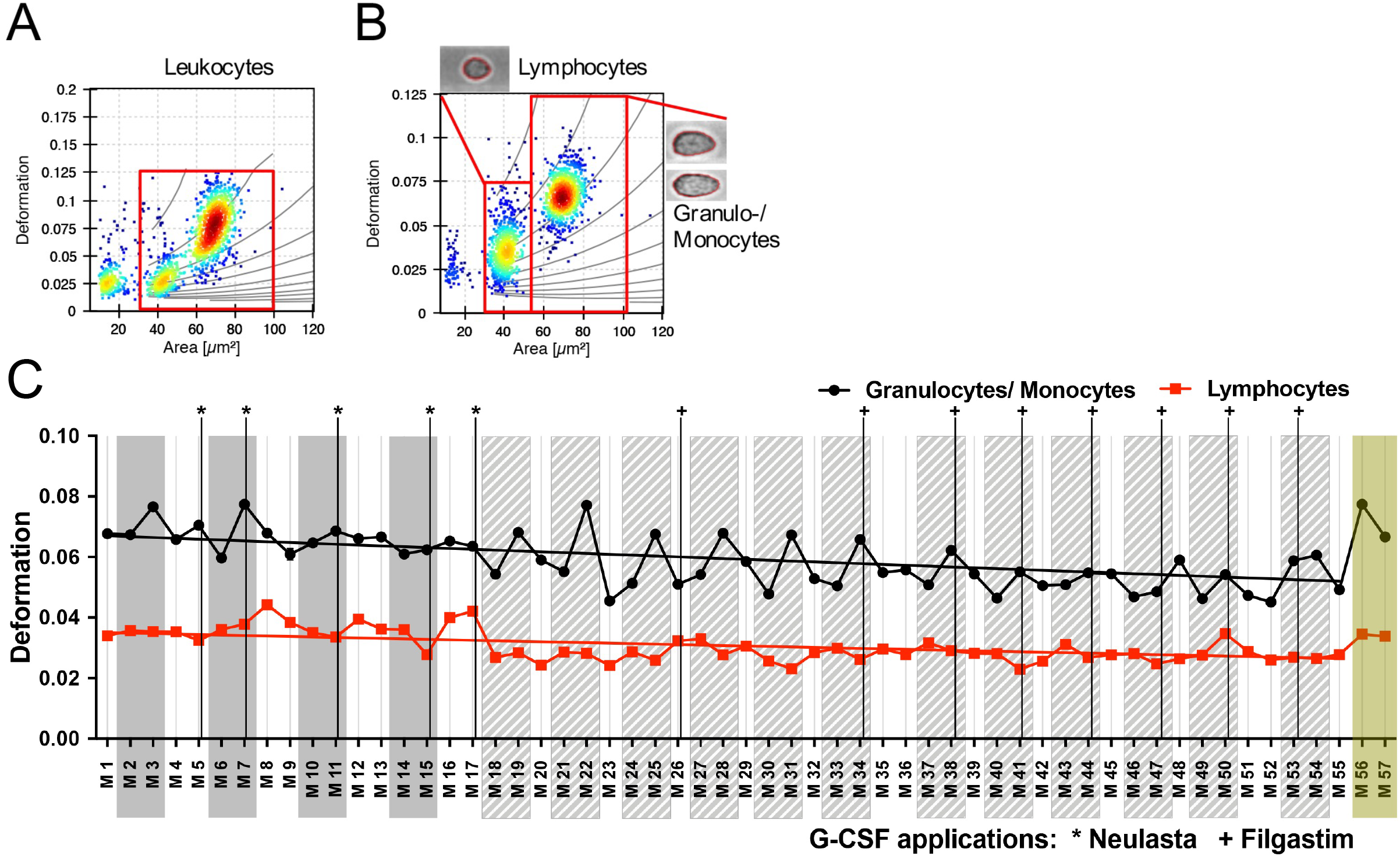
*(A+B)* Gating strategy to separate lymphocytes from granulo-/monocytes of representative scatter plots of deformation versus area. Gates: area ratio from 1 to 1.05; Lymphocyte gating area 30-50 µm^2^ and gating deformation 0-0.075. Granulo-/ monocyte gating for area 50-100 µm^2^ and gating for deformation 0-0.125 (A) Dots show all cells that were measured. (B) Images show an example of a cell either from the lymphocyte gate or from the granulo/monocyte gate. Thin, curved lines in the background are isoelasticity lines that indicate that indicate regions of equal apparent Young’s modulus (YM). *(C)* 57 RT-DC blood measurements (M_1-57_) were performed during the course of this study as indicated by rectangles. During the chemotherapy treatment, measurements were done immediately before and after every treatment cycle as well as three days after treatment (white zones between the grey boxes). Additionally, measurements M36 and M52 were carried out one day later because the patient’s therapy schedule had to be postponed by one day for health reasons. Grey boxes indicate epirubicin/ cyclophosphamide (E/C) treatment (seven weeks, biweekly treatment) followed by 12 cycles of paclitaxel treatment (Pax) (twelve weeks, weekly treatment) striped light grey boxes. Administration of G-CSF (*Neulasta or ^+^Filgastrim) is shown by vertical lines. Deformation values of granulo-/ monocytes (interconnected black dots) and lymphocytes (interconnected red squares) obtained from RT-DC measurements before onset of the chemotherapeutic intervention, throughout the therapy and in remission. Dots (Granulo-/ monocytes) and squares (Lymphocytes) are the median values. The line is only a graphical aid for the time-dependent changes and can be seen as a visual guide.

