## Supplement table 1-2 for "Impact of chemotherapy on leukocyte stiffness -A longitudinal RT-DC study in a breast cancer patient"

*Supplement Table 1:* Median values, standard deviations (SDs) and standard error (SEMs) for apparent Young’s modulus (YM), area and deformation for M1-M57 for granulo-/ monocytes.

|  | YM [kPa] | SEM  YM [kPa] | SD  YM [kPa] | Area [µm²] | SEM  Area [µm²] | SD  Area [µm²] | Deformation | SEM  Deformation | SD  Deformation |
| --- | --- | --- | --- | --- | --- | --- | --- | --- | --- |
| M 1 | 0.848 | 0.00248 | 0.07307 | 69.8 | 0.17755 | 5.23693 | 0.0677 | 0.00037 | 0.01098 |
| M 2 | 0.854 | 0.01559 | 0.36866 | 70.5 | 0.23968 | 5.66670 | 0.0674 | 0.00055 | 0.01307 |
| M 3 | 0.828 | 0.00632 | 0.18202 | 73.5 | 0.18071 | 5.20319 | 0.0766 | 0.00035 | 0.01013 |
| M 4 | 0.868 | 0.00186 | 0.06563 | 71.3 | 0.15554 | 5.47925 | 0.0658 | 0.00029 | 0.01019 |
| M 5 | 0.883 | 0.00422 | 0.06870 | 77.4 | 0.38091 | 6.20082 | 0.0705 | 0.00069 | 0.01118 |
| M 6 | 0.906 | 0.02125 | 0.32779 | 70.0 | 0.41947 | 6.47132 | 0.0597 | 0.00120 | 0.01856 |
| M 7 | 0.841 | 0.00462 | 0.12006 | 76.2 | 0.23809 | 6.19043 | 0.0774 | 0.00050 | 0.01305 |
| M 8 | 0.878 | 0.00197 | 0.09346 | 73.8 | 0.14474 | 6.87317 | 0.0679 | 0.00032 | 0.01509 |
| M 9 | 0.928 | 0.08762 | 0.93555 | 77.0 | 1.04759 | 11.18520 | 0.0609 | 0.00191 | 0.02042 |
| M 10 | 0.864 | 0.00435 | 0.10892 | 69.6 | 0.22210 | 5.55698 | 0.0647 | 0.00045 | 0.01134 |
| M 11 | 0.856 | 0.00213 | 0.08140 | 71.8 | 0.14767 | 5.65579 | 0.0686 | 0.00028 | 0.01069 |
| M 12 | 0.876 | 0.00120 | 0.07109 | 72.6 | 0.10597 | 6.29363 | 0.0661 | 0.00019 | 0.01151 |
| M 13 | 0.924 | 0.00420 | 0.08485 | 80.1 | 0.33774 | 6.83039 | 0.0666 | 0.00060 | 0.01220 |
| M 14 | 0.886 | 0.00599 | 0.13785 | 68.9 | 0.25542 | 5.87460 | 0.0610 | 0.00056 | 0.01287 |
| M 15 | 0.883 | 0.00911 | 0.30722 | 70.0 | 0.16581 | 5.59095 | 0.0624 | 0.00031 | 0.01061 |
| M 16 | 0.869 | 0.00072 | 0.07057 | 70.8 | 0.06103 | 5.94121 | 0.0653 | 0.00011 | 0.01117 |
| M 17 | 0.935 | 0.01139 | 0.12263 | 79.1 | 0.83655 | 9.00988 | 0.0635 | 0.00143 | 0.01535 |
| M 18 | 0.926 | 0.00379 | 0.12023 | 68.4 | 0.18013 | 5.70747 | 0.0543 | 0.00053 | 0.01665 |
| M 19 | 0.799 | 0.00282 | 0.10765 | 63.1 | 0.14871 | 5.69742 | 0.0681 | 0.00052 | 0.01992 |
| M 20 | 0.864 | 0.00393 | 0.08196 | 64.9 | 0.22419 | 4.67585 | 0.0590 | 0.00054 | 0.01133 |
| M 21 | 0.912 | 0.00380 | 0.12037 | 67.0 | 0.16341 | 5.17012 | 0.0552 | 0.00035 | 0.01117 |
| M 22 | 0.796 | 0.00387 | 0.11352 | 68.4 | 0.19087 | 5.59736 | 0.0771 | 0.00069 | 0.02032 |
| M 23 | 0.992 | 0.00488 | 0.15084 | 64.9 | 0.17059 | 5.27735 | 0.0455 | 0.00037 | 0.01139 |
| M 24 | 0.957 | 0.00454 | 0.12754 | 68.4 | 0.19482 | 5.47223 | 0.0513 | 0.00039 | 0.01087 |
| M 25 | 0.833 | 0.00549 | 0.12512 | 68.3 | 0.26767 | 6.09802 | 0.0675 | 0.00090 | 0.02044 |
| M 26 | 0.933 | 0.00642 | 0.10604 | 65.5 | 0.31149 | 5.14664 | 0.0510 | 0.00060 | 0.00987 |
| M 27 | 0.929 | 0.00211 | 0.08874 | 68.3 | 0.12438 | 5.24320 | 0.0542 | 0.00024 | 0.01000 |
| M 28 | 0.845 | 0.00492 | 0.13921 | 69.5 | 0.20288 | 5.74197 | 0.0678 | 0.00059 | 0.01672 |
| M 29 | 0.918 | 0.00545 | 0.10270 | 71.2 | 0.30452 | 5.73756 | 0.0585 | 0.00070 | 0.01316 |
| M 30 | 0.962 | 0.00818 | 0.13191 | 64.3 | 0.30835 | 4.97194 | 0.0478 | 0.00066 | 0.01058 |
| M 31 | 0.819 | 0.00872 | 0.14361 | 65.3 | 0.35756 | 5.88619 | 0.0673 | 0.00132 | 0.02171 |
| M 32 | 0.934 | 0.00645 | 0.12152 | 66.8 | 0.30765 | 5.79650 | 0.0528 | 0.00066 | 0.01252 |
| M 33 | 0.970 | 0.00496 | 0.11214 | 68.1 | 0.25748 | 5.82031 | 0.0505 | 0.00045 | 0.01007 |
| M 34 | 0.847 | 0.02325 | 0.50304 | 68.6 | 0.30439 | 6.60946 | 0.0658 | 0.00096 | 0.02096 |
| M 35 | 0.941 | 0.00279 | 0.11641 | 70.5 | 0.14121 | 5.89018 | 0.0549 | 0.00023 | 0.00972 |
| M 36 | 0.940 | 0.00358 | 0.10822 | 71.2 | 0.22839 | 6.90110 | 0.0558 | 0.00044 | 0.01336 |
| M 37 | 0.973 | 0.00985 | 0.17230 | 69.5 | 0.38681 | 6.76644 | 0.0508 | 0.00066 | 0.01154 |
| M 38 | 0.876 | 0.02126 | 0.39425 | 69.2 | 0.40091 | 7.44842 | 0.0622 | 0.00101 | 0.01913 |
| M 39 | 0.932 | 0.01092 | 0.21832 | 66.6 | 0.29609 | 5.92173 | 0.0544 | 0.00061 | 0.01216 |
| M 40 | 1.000 | 0.00372 | 0.10738 | 67.6 | 0.19007 | 5.49238 | 0.0464 | 0.00033 | 0.00963 |
| M 41 | 0.915 | 0.01414 | 0.32237 | 68.1 | 0.28770 | 6.56058 | 0.0551 | 0.00086 | 0.01958 |
| M 42 | 0.944 | 0.00674 | 0.12015 | 65.8 | 0.30929 | 5.51545 | 0.0506 | 0.00073 | 0.01302 |
| M 43 | 0.960 | 0.00411 | 0.13415 | 67.9 | 0.17533 | 5.73313 | 0.0509 | 0.00033 | 0.01073 |
| M 44 | 0.926 | 0.01086 | 0.26971 | 67.5 | 0.24297 | 6.03754 | 0.0548 | 0.00067 | 0.01673 |
| M 45 | 0.923 | 0.01937 | 0.34595 | 68.1 | 0.29924 | 5.33674 | 0.0545 | 0.00070 | 0.01280 |
| M 46 | 0.992 | 0.00615 | 0.16864 | 67.3 | 0.21247 | 5.83043 | 0.0469 | 0.00053 | 0.01454 |
| M 47 | 0.973 | 0.01750 | 0.41152 | 67.4 | 0.26657 | 6.26863 | 0.0485 | 0.00072 | 0.01691 |
| M 48 | 0.906 | 0.04194 | 0.54038 | 68.9 | 0.48145 | 6.18520 | 0.0590 | 0.00117 | 0.01560 |
| M 49 | 1.000 | 0.00425 | 0.12599 | 67.2 | 0.19399 | 5.75473 | 0.0463 | 0.00033 | 0.00971 |
| M 50 | 0.940 | 0.00395 | 0.13366 | 69.2 | 0.17989 | 6.08170 | 0.0542 | 0.00041 | 0.01382 |
| M 51 | 1.010 | 0.03590 | 0.59422 | 68.6 | 0.34794 | 5.85811 | 0.0473 | 0.00079 | 0.01322 |
| M 52 | 1.020 | 0.00254 | 0.12558 | 67.7 | 0.11187 | 5.54180 | 0.0451 | 0.00021 | 0.01032 |
| M 53 | 0.902 | 0.00551 | 0.10243 | 69.9 | 0.31052 | 5.77593 | 0.0588 | 0.00070 | 0.01294 |
| M 54 | 0.876 | 0.00513 | 0.14829 | 68.3 | 0.20769 | 6.00881 | 0.0606 | 0.00061 | 0.01753 |
| M 55 | 0.972 | 0.00760 | 0.12446 | 68.0 | 0.31072 | 5.08675 | 0.0492 | 0.00070 | 0.01151 |
| M 56 | 0.803 | 0.00301 | 0.06860 | 69.4 | 0.23759 | 5.41278 | 0.0774 | 0.00051 | 0.01173 |
| M 57 | 0.843 | 0.00581 | 0.19480 | 67.8 | 0.16011 | 5.38280 | 0.0666 | 0.00036 | 0.01204 |

*Supplement Table 2:* Median values, standard deviations (SDs) and standard error (SEMs) for apparent Young’s modulus (YM), area and deformation for M1-M57 for lymphocytes.

|  | Median YM [kPa] | SEM  YM [kPa] | SD  YM [kPa] | Area [µm²] | SEM  Area [µm²] | SD Area [µm²] | Deformation | SEM  Deformation | SD  Deformation |
| --- | --- | --- | --- | --- | --- | --- | --- | --- | --- |
| M 1 | 0.924 | 0.01588 | 0.38169 | 39.7 | 0.14066 | 3.78091 | 0.03400 | 0.00043 | 0.01041 |
| M 2 | 0.905 | 0.01121 | 0.24113 | 40.7 | 0.15261 | 3.63327 | 0.03570 | 0.00047 | 0.01013 |
| M 3 | 0.894 | 0.02081 | 0.23083 | 40.4 | 0.32039 | 3.86517 | 0.03540 | 0.00102 | 0.01148 |
| M 4 | 0.891 | 0.01354 | 0.33260 | 38.5 | 0.13453 | 3.60594 | 0.03530 | 0.00047 | 0.01133 |
| M 5 | 0.964 | 0.01829 | 0.30545 | 40.1 | 0.20911 | 3.93237 | 0.03250 | 0.00065 | 0.01070 |
| M 6 | 0.904 | 0.03571 | 0.55324 | 42.4 | 0.23311 | 3.97743 | 0.03610 | 0.00079 | 0.01272 |
| M 7 | 0.897 | 0.08254 | 0.75197 | 42.9 | 0.39096 | 4.26828 | 0.03780 | 0.00139 | 0.01293 |
| M 8 | 0.813 | 0.02063 | 0.33009 | 42.6 | 0.23125 | 3.80050 | 0.04420 | 0.00066 | 0.01050 |
| M 9 | 0.889 | 0.01530 | 0.26280 | 43.4 | 0.23003 | 4.32235 | 0.03840 | 0.00064 | 0.01127 |
| M 10 | 0.933 | 0.01606 | 0.24088 | 43.1 | 0.22953 | 3.91305 | 0.03500 | 0.00063 | 0.00958 |
| M 11 | 0.943 | 0.04296 | 0.44855 | 43.5 | 0.34400 | 4.16241 | 0.03360 | 0.00117 | 0.01251 |
| M 12 | 0.858 | 0.01416 | 0.22613 | 42.5 | 0.24041 | 4.28534 | 0.03950 | 0.00068 | 0.01121 |
| M 13 | 0.912 | 0.02213 | 0.33262 | 41.7 | 0.25737 | 4.23639 | 0.03620 | 0.00075 | 0.01139 |
| M 14 | 0.918 | 0.03234 | 0.46420 | 43.5 | 0.25480 | 4.20175 | 0.03600 | 0.00079 | 0.01156 |
| M 15 | 1.100 | 0.07214 | 0.33057 | 42.8 | 0.78223 | 5.31753 | 0.02780 | 0.00152 | 0.00942 |
| M 16 | 0.852 | 0.01643 | 0.32162 | 41.5 | 0.18797 | 3.93049 | 0.04000 | 0.00055 | 0.01095 |
| M 17 | 0.825 | 0.01702 | 0.19993 | 42.5 | 0.31215 | 4.12896 | 0.04210 | 0.00087 | 0.01009 |
| M 18 | 1.120 | 0.03295 | 0.51356 | 42.4 | 0.22653 | 3.78083 | 0.02680 | 0.00068 | 0.01063 |
| M 19 | 1.040 | 0.07041 | 0.60974 | 37.6 | 0.40013 | 4.19156 | 0.02840 | 0.00119 | 0.01130 |
| M 20 | 1.160 | 0.04787 | 0.40051 | 40.2 | 0.40187 | 3.86488 | 0.02430 | 0.00108 | 0.00902 |
| M 21 | 1.060 | 0.01703 | 0.37928 | 42.1 | 0.15931 | 3.86002 | 0.02860 | 0.00046 | 0.01047 |
| M 22 | 1.070 | 0.01804 | 0.29759 | 42.9 | 0.21848 | 3.99740 | 0.02820 | 0.00060 | 0.00995 |
| M 23 | 1.210 | 0.04705 | 0.61883 | 43.9 | 0.26240 | 3.82143 | 0.02420 | 0.00075 | 0.00989 |
| M 24 | 1.070 | 0.01728 | 0.35023 | 42.5 | 0.16355 | 3.54893 | 0.02870 | 0.00049 | 0.01014 |
| M 25 | 1.160 | 0.02398 | 0.33741 | 42.1 | 0.22397 | 3.26718 | 0.02590 | 0.00063 | 0.00886 |
| M 26 | 0.999 | 0.03952 | 0.39911 | 43.0 | 0.34396 | 4.13294 | 0.03230 | 0.00104 | 0.01100 |
| M 27 | 0.970 | 0.02728 | 0.52677 | 42.1 | 0.17612 | 3.58648 | 0.03300 | 0.00051 | 0.00994 |
| M 28 | 1.090 | 0.04272 | 0.42295 | 42.4 | 0.37146 | 4.64286 | 0.02770 | 0.00096 | 0.01087 |
| M 29 | 1.020 | 0.04845 | 0.56295 | 42.6 | 0.29408 | 3.57966 | 0.03060 | 0.00092 | 0.01076 |
| M 30 | 1.150 | 0.07564 | 0.84566 | 41.2 | 0.29125 | 3.47545 | 0.02570 | 0.00085 | 0.00954 |
| M 31 | 1.230 | 0.03044 | 0.45355 | 40.7 | 0.21862 | 3.69178 | 0.02310 | 0.00059 | 0.00926 |
| M 32 | 1.090 | 0.03338 | 0.38785 | 43.0 | 0.26960 | 3.48326 | 0.02840 | 0.00086 | 0.01015 |
| M 33 | 1.030 | 0.01618 | 0.26342 | 42.4 | 0.19956 | 3.39650 | 0.02990 | 0.00059 | 0.00952 |
| M 34 | 1.150 | 0.07414 | 1.19998 | 43.0 | 0.21111 | 3.93268 | 0.02610 | 0.00061 | 0.01013 |
| M 35 | 1.040 | 0.03414 | 0.44125 | 42.8 | 0.26538 | 3.49306 | 0.02960 | 0.00080 | 0.01054 |
| M 36 | 1.070 | 0.01990 | 0.28771 | 42.1 | 0.23621 | 3.57044 | 0.02780 | 0.00065 | 0.00931 |
| M 37 | 1.020 | 0.02191 | 0.30825 | 42.8 | 0.23847 | 3.56412 | 0.03160 | 0.00070 | 0.00981 |
| M 38 | 1.040 | 0.02930 | 0.43656 | 42.1 | 0.21487 | 3.80402 | 0.02910 | 0.00063 | 0.01009 |
| M 39 | 1.090 | 0.03170 | 0.40349 | 42.8 | 0.27019 | 3.51171 | 0.02820 | 0.00081 | 0.01053 |
| M 40 | 1.080 | 0.07213 | 1.11744 | 42.0 | 0.22355 | 3.83121 | 0.02810 | 0.00068 | 0.01062 |
| M 41 | 1.230 | 0.02812 | 0.41329 | 40.7 | 0.23287 | 3.77602 | 0.02300 | 0.00054 | 0.00878 |
| M 42 | 1.140 | 0.02768 | 0.39344 | 41.3 | 0.22975 | 3.64400 | 0.02560 | 0.00073 | 0.01021 |
| M 43 | 1.010 | 0.02635 | 0.44711 | 41.9 | 0.20980 | 4.35221 | 0.03120 | 0.00062 | 0.01103 |
| M 44 | 1.130 | 0.01629 | 0.26664 | 42.7 | 0.19887 | 3.78137 | 0.02680 | 0.00049 | 0.00842 |
| M 45 | 1.080 | 0.03002 | 0.49508 | 42.0 | 0.21841 | 3.81581 | 0.02770 | 0.00052 | 0.00877 |
| M 46 | 1.070 | 0.01854 | 0.31295 | 41.6 | 0.19610 | 3.52621 | 0.02810 | 0.00056 | 0.00954 |
| M 47 | 1.190 | 0.02363 | 0.37737 | 41.9 | 0.20699 | 3.73735 | 0.02470 | 0.00057 | 0.00957 |
| M 48 | 1.100 | 0.03353 | 0.36884 | 40.9 | 0.29640 | 3.75656 | 0.02640 | 0.00095 | 0.01097 |
| M 49 | 1.110 | 0.01782 | 0.29282 | 41.6 | 0.21140 | 3.90294 | 0.02760 | 0.00055 | 0.00963 |
| M 50 | 0.943 | 0.01173 | 0.23637 | 42.8 | 0.17758 | 3.76434 | 0.03470 | 0.00050 | 0.01019 |
| M 51 | 1.060 | 0.03020 | 0.41622 | 42.1 | 0.24795 | 4.08576 | 0.02880 | 0.00069 | 0.01001 |
| M 52 | 1.120 | 0.02209 | 0.32084 | 40.9 | 0.24269 | 3.79895 | 0.02600 | 0.00058 | 0.00872 |
| M 53 | 1.100 | 0.03337 | 0.50062 | 41.0 | 0.22143 | 3.43831 | 0.02690 | 0.00064 | 0.00952 |
| M 54 | 1.120 | 0.02948 | 0.62320 | 39.4 | 0.17116 | 4.80156 | 0.02650 | 0.00046 | 0.01061 |
| M 55 | 1.080 | 0.06999 | 0.97731 | 41.1 | 0.24384 | 3.66741 | 0.02780 | 0.00070 | 0.01012 |
| M 56 | 0.922 | 0.01445 | 0.23473 | 41.3 | 0.22706 | 4.06116 | 0.03450 | 0.00065 | 0.01053 |
| M 57 | 0.935 | 0.01821 | 0.32622 | 40.3 | 0.20548 | 4.09565 | 0.03380 | 0.00059 | 0.01058 |
